# Analyzing editosome function in high-throughput

**DOI:** 10.1101/2020.05.15.096370

**Authors:** Cristian Del Campo, Wolf-Matthias Leeder, Paul Reißig, H. Ulrich Göringer

## Abstract

Mitochondrial gene expression in African trypanosomes and other trypanosomatid pathogens requires a U-nucleotide specific insertion/deletion-type RNA-editing reaction. The process is catalyzed by a macromolecular protein complex known as the editosome. Editosomes are restricted to the trypanosomatid clade and since editing is essential for the parasites, the protein complex represents a near perfect target for drug intervention strategies. Here we report the development of an improved *in vitro* assay to monitor editosome function. The test system utilizes fluorophore-labeled substrate RNAs to analyze the processing reaction by automated, high-throughput capillary electrophoresis (CE) in combination with a laser-induced fluorescence (LIF) readout. We optimized the assay for high-throughput screening (HTS)-experiments and devised a multiplex fluorophore-labeling regime to scrutinize the U-insertion/U-deletion reaction simultaneously. The assay is robust, it requires only nanogram amounts of materials and it meets all performance criteria for HTS-methods. As such the test system should be helpful in the search for trypanosome-specific pharmaceuticals.

## Introduction

Human African trypanosomiasis (HAT), also known as sleeping sickness, is a neglected tropical disease. Despite a decreasing number of new infections in recent years, still 70 million people in 36 African countries are at risk of becoming infected (Kennedy, 2019). In response to the declining incidence, the World Health Organization (WHO) has targeted HAT for elimination as a public health problem by 2020 (Barrett, 2018). However, a similar situation was already achieved in the 1960s, which was followed by a reduction in surveillance and control activities and as a consequence the disease resurfaced again in the 1990s (Brun et al., 2010). This advocates that efforts to develop trypanosome-specific diagnostics and therapeutics should not be suspended (Burri, 2020). Five drugs are currently used in the treatment of HAT (Büscher et al., 2017). All of them are toxic to different degrees (Kennedy, 2019). In addition, they suffer from a multitude of complications including parenteral administration (Gilbert, 2014), poor efficacy, clinical side effects and increasing levels of resistance (Baker et al., 2013; Barrett et al., 2011; Fairlamb and Horn, 2018). As a consequence, a novel and improved HAT-therapeutic would be of great value (Field et al., 2017). Causative agent of HAT is *Trypanosoma brucei*, an extracellular, single-cell parasite. The organism proliferates in the blood and lymph fluid and is capable of crossing the blood-brain barrier where it induces a wide spectrum of neurological symptoms (Büscher et al., 2017). Left untreated, progressive mental deterioration leads to coma and death (Jamonneau et al., 2012).

Mitochondrial gene expression in trypanosomes and related pathogens requires an RNA-editing reaction in which sequence-deficient, non-translatable primary transcripts are converted into translatable mRNAs by the side-specific insertion and deletion of exclusively U-nucleotides (Benne et al. 1986; reviewed in Aphasizheva et al., 2020). The reaction pathway is catalyzed by a unique, high molecular mass multienzyme complex known as the editosome (Pollard et al., 1992; Golas et al., 2009; reviewed in Göringer, 2012). Editosomes execute the processing reaction in a cascade of enzyme-mediated steps, which include RNA-chaperone, endo- and exonuclease, terminal uridylyl transferase (TUTase) and RNA-ligase activities. Furthermore, the process is mediated by small, non-coding RNAs, termed guide (g)RNAs. They act as templates and direct the U-insertion/deletion by anti-parallel base pairing (Blum et al., 1990; Hermann et al., 1997). Several features make the editosome a prime drug target. First, RNA-editing is an essential pathway in trypanosomes. Second, the editosome is uniquely present in the parasite, with no corresponding enzyme complex in the host. Third, the reaction cycle involves multiple enzyme reactions, all representing potential drug-interference points, and fourth, the high molecular mass protein complex (0.8MDa) offers a large drug-binding landscape (Golas et al., 2009). Despite these favorable features, only very few RNA-editing inhibitors have been identified (Liang and Connell, 2010; Moshiri and Salavati, 2010; Katari et al., 2013, Leeder et al., 2015; Zimmermann et al., 2016). Moreover, only three studies have systematically searched for RNA-editing inhibiting compounds, which primarily is due to the lack of a robust, high-throughput compatible assay system. Available formats include an electrochemiluminescent RNA aptamer-based test (Liang and Connell, 2009), a hammerhead ribozyme (HHR)-driven reporter assay connected to Förster resonance energy transfer (FRET)-detection (Moshiri et al., 2014) and a second FRET-based system to detect RNA-ligase 1-specific inhibitors (Zimmermann et al., 2016). Despite the fact that the described assays successfully identified RNA editing-specific inhibitors, the different formats also pose limitations. These range from relying on indirect measuring principles to focusing exclusively on one step of the reaction cycle. Most importantly however, none of the methods is able to provide a complete and quantitative analysis of all editosome-mediated catalytic conversion steps.

Here we present a new, high-throughput RNA-editing screening assay. The test system is a modified version of the established *in vitro* RNA-editing assays (Seiwert and Stuart, 1994, Kable et al., 1996; Stuart et al., 1998), but instead of using radioactively labeled substrate-RNAs it utilizes fluorophore-derivatized RNA-substrates (Fig. 1). This enables the usage of automated, highly parallelized capillary electrophoresis (CE) instruments coupled to laser-induced fluorescence (LIF) detection systems. We optimized the work flow of the assay by adjusting the material quantities and reaction volume to multiwell-plate formats, by increasing the RNase-stability of the substrate-RNAs and by devising a multiplex fluorophore-labeling regime to scrutinize the U-insertion and U-deletion reaction in one reaction vial using chemically identical U-insertion/U-deletion substrate RNAs. The assay is robust, it satisfies all signal-to-noise criteria for high-throughput screening methods and it is able to derive quantitative data for every reaction step of the catalytic conversion. To validate the assay, we surveyed UTP-analogs for their aptitude to inhibit the U-insertion reaction.

**Figure 1.**
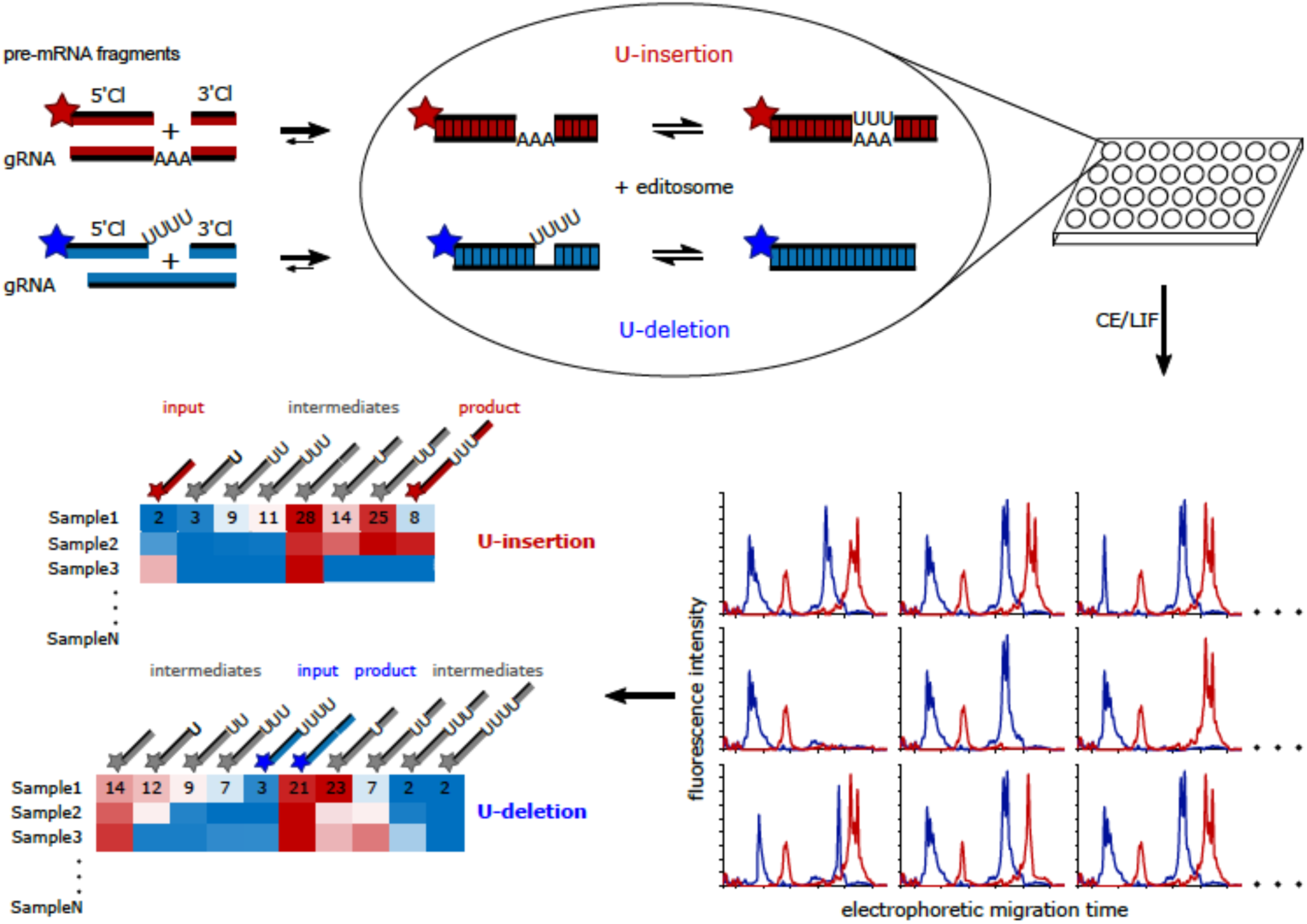
Work flow of the fluorescence-based insertion/deletion editing (FIDE)-assay. The test system uses short, synthetic oligoribonucleotides to mimic the pre-edited mRNAs and gRNAs of the editing reaction. Two sets of RNA-oligonucleotides are synthesized to monitor the U-insertion and U-deletion reaction separately. Pre-mRNA molecules are introduced as pre-cleaved RNAs (5’-cleavage (Cl)-fragment, 3’-cleavage (Cl)-fragment) to sidestep the rate-limiting endonucleolytic cleavage reaction. Fluorophore-derivatized versions of the pre-mRNA oligonucleotides (stars in red and blue) are annealed to their base-complementary gRNAs to form trimolecular pre-mRNA/gRNA hybrid RNAs, which represent the substrates of the editing reaction. Upon incubation with editosomes the pre-mRNAs are edited by the insertion of 3 U-nt or the deletion of 4 U’s. Reactions are performed in multiwell plates and are subsequently analyzed by automated capillary electrophoresis (CE) coupled to laser-induced fluorescence (LIF) detection. Using different fluorophores for the U-insertion and U-deletion, both reactions can be monitored simultaneously. The resulting electropherograms are peak integrated, facilitating a quantitative side-by-side comparison of every product and intermediate of the two reactions. The usage of multi-capillary CE-instruments provides a highly parallelized, high-throughput environment, in which hundreds of samples can be analyzed at the same time. As such the assay is well-suited for drug-screening purposes.

## Materials and Methods

### Chemical RNA-synthesis and postsynthetic processing

RNA-oligonucleotides were synthesized by solid-phase synthesis on controlled pore glass (CPG)-beads (50nmol synthesis scale) using 2′-O-(tert-butyl)dimethylsilyl (TBDMS)-protected phosphoramidites. 5-Carboxytetramethyl-rhodamine (TAMRA, λ_ex_ 546nm, λ_em_ 579nm), 6-hexachloro-fluorescein (HEX, λ_ex_ 535nm, λ_em_ 556nm) and 6-carboxy-fluorescein (FAM, λ_ex_ 492nm, λ_em_ 517nm) were chosen as fluorophore substituents and were introduced post-synthetically either at the 5’- or 3’-ends of the different oligoribonucleotides. For that the RNA-oligos were synthesized to contain either a 5’- or 3’-terminal C6-aminolinker. The primary amino group was then conjugated to carboxyl functional groups of FAM, TAMRA and HEX using an EDC (1-Ethyl-3-(3-dimethylaminopropyl)carbodiimide)-mediated coupling reaction. As additional modifications, some RNAs were synthesized to include up to 6 phosphorothioate (PS) backbone modifications to increase their RNase-stability. Similarly, some of the 3’-pre-mRNA oligonucleotides were 3’-end modified by introducing a hexamethylene-amino-linker. Base-protecting groups were removed at mild conditions using NH_4_OH/EtOH (3:1) at RT and 2’-silyl protecting groups were removed using neat triethylamine trihydrofluoride. All RNA-oligonucleotides were HPLC-purified, analyzed by mass-spectrometry and further scrutinized in denaturing polyacrylamide gels (suppl. Fig. S1). RNA-concentration were derived from UV-absorbency measurements at 260nm (A_260_) using the molar extinction coefficients (ε in L/mol x cm) listed below. Oligoribonucleotides representing 3’-pre-mRNA fragments were enzymatically 5’-phosphorylated using T4-polynucleotide kinase (T4-PNK) and ATP using standard conditions. The following RNA-oligonucleotides were synthesized:

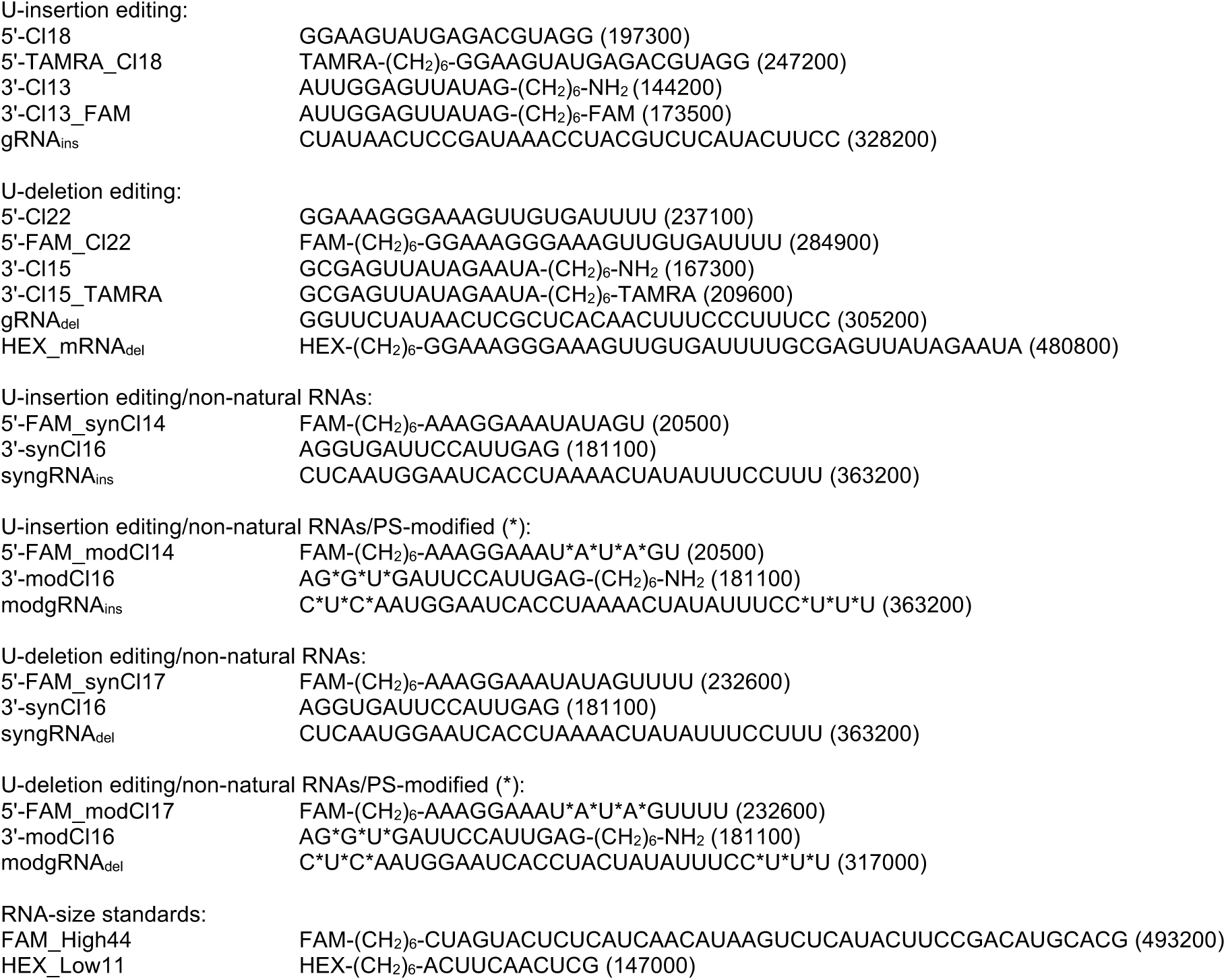

### Formation of pre-mRNA/gRNA-hybrid RNAs

Trimolecular pre-mRNA/gRNA-hybrid molecules were generated by hybridization of the individual 5’- and 3’-pre-mRNA oligoribonucleotides to base complementary gRNA molecules. For that, equimolar amounts (4pmol) of the three RNA-oligonucleotides were combined in a final volume of 0.1mL 10mM Tris/HCl, 1mM Na_2_EDTA (TE) pH7.5 followed by denaturation at 75°C for 2min. Denatured oligoribonucleotides were annealed by cooling samples to 27°C at a rate of 0.08°C/s. Formation of the pre-mRNA/gRNA-hybrid RNAs was verified by electrophoresis in native 15% (w/v) polyacrylamide gels followed by staining with Toludine blue O. Stained gels were densitometrically analyzed.

### Editosome purification

Editosomes were purified from insect-stage African trypanosomes (*Trypanosoma brucei*), which were grown at 27°C in SDM-79 medium (Brun and Schönenberger, 1979) in the presence of 10% (v/v) fetal calf serum. The complexes were either purified by tandem-affinity purification (TAP) (Rigaut et al., 1999; Golas et al., 2009; Gerace and Moazed, 2015) using transgenic *T. brucei* cell lines that conditionally express TAP-tagged versions of editosomal proteins or from mitochondrial detergent extracts of wildtype trypanosomes (strain Lister 427) as described in Böhm et al., 2012. The following transgenic parasite strains were used: *T. brucei* 29-13 KREPA4/TAP (Kala et al., 2012), *T. brucei* 29-13 KREPA3/TAP (Golas et al., 2009) and *T. brucei* 29-13 KRET2/TAP (Ernst et al., 2003; Ringpis et al., 2010). Typically about 6×10^11^ parasites were disrupted at isotonic, near-native conditions and cell lysates were processed using IgG- and calmodulin-affinity chromatography resins. Protein concentrations were determined by Bradford dye binding and the protein inventory of the different isolates was analyzed by tandem mass spectrometry (nanoLC-MS/MS).

### FIDE - Fluorescence-based Insertion/Deletion Editing

RNA-editing reactions were assembled in a 30μL volume in editing buffer (EB: 20mM HEPES pH7.5, 10mM MgCl_2_, 30mM KCl) supplemented with 0.5mM DTT and 0.5mM ATP. For the U-insertion assay, additional 0.1mM UTP or analogs of UTP were added. Reactions contained 0.4pmol pre-annealed pre-mRNA/gRNA hybrid RNA and 0.2-0.4pmoles of editosomes. Samples were incubated at 27°C for 3h after which 15fmol of two size-standard oligoribonucleotides (FAM_High44 and HEX_Low11) were added. Reactions were stopped by phenol/chloroform extraction and the processed RNAs were EtOH precipitated. Samples were immediately centrifuged for 30min at 13000rpm (4°C) and RNA pellets were washed (70% (v/v) EtOH), dried *in vacuo* and resolved in 100μL of Hi-Di™ formamide. Samples were heat denatured at 95°C for 2min, snap-cooled and separated by capillary electrophoresis (CE) at 12kV for 25min using a POP-6 polymer (Thermo Scientific). “One-pot” U-insertion/U-deletion reactions were assembled by annealing of the U-insertion and U-deletion pre-mRNA/gRNA-hybrid RNAs in separate reaction vials. Equimolar amounts (0.2pmol) of each hybrid RNA were then combined with 0.2-0.4pmol of editosomes in EB in the presence of 0.5mM ATP, 0.1mM UTP and 0.5mM DTT in a final volume of 30µL. All incubation and processing steps of the one-pot samples were performed as above. Down-scaled U-insertion and U-deletion assays were performed with 10fmol of pre-mRNA/gRNA hybrid RNAs and 2.5fmoles of editosomes in a volume of 4µL. The reduced amounts of material made it possible to skip the phenol extraction and EtOH-precipitation steps of the work flow and enabled a direct sample application onto the CE-system. We also confirmed that the FIDE-assay can be performed in a non-precleaved fashion. For that we used the pre-mRNA mimicking oligoribonucleotide HEX_mRNA_del_ in conjunction with gRNA_del_.

### Data processing

Raw data from the CE-runs (relative fluorescence units (RFU) and electrophoretic migration times in seconds) were converted into tab-delimited text files and imported either into OriginPro v8.5 (OriginLab Corporation) or MultiGauge v3.0 (Fuji Photo Film, Co.). The quality of the CE-separations was assessed by evaluating the signal-to-noise (S/N) ratio as S/N=(µ_signal_/µ_noise_)/∂_noise_ (µ=mean; ∂=standard deviation). The data were baseline corrected and individual peaks were assigned by comparison to the unedited peak of a mock reaction using the electrophoretic elution times of two 11nt- and 44nt-long calibration oligoribonucleotides (FAM_High44 and HEX_Low11) as reference points. Peaks were integrated using a Riemann sum approximation to derive RNA-editing activity values (EA) defined as the ratio of the peak area (A) of the fully edited product (A_FE_) *versus* the sum of all peaks *i.e*. the unedited (A_UE_), partially edited (A_PE_) and fully edited (A_FE_): EA=A_FE_ / ∑(A_UE,_ A_PE_, A_FE_). EA-values divided by the mass of editosomes represent the specific RNA-editing activity of the editosome preparation in EA/ng. By computing the ratio between the area of each peak and the total integrated area, the fraction of each reaction intermediate was calculated. In the case of “one-pot” RNA-editing assays, raw data from the electropherograms were first imported into ShapeFinder (Vasa et al. 2008) for baseline correction and matrixing in order to correct for the unique contribution of each fluorophore to the signal intensity in each fluorescence channel. The quality of the HTS-experiment was assessed by calculating the mean screening window coefficient (Z’) as Z’=1-(3∂_control+_ + 3∂_control-_) / |µ_control+_-µ_control-_| (µ=mean, ∂=standard deviation, control+ = positive control, control-= negative control) (Zhang et al., 1999).

### UV-hyperchromicity measurements

Absorbance *versus* temperature profiles (melting curves) of the different pre-mRNA/gRNA hybrid RNAs were recorded at 260nm (A_260_) using a thermoelectrically controlled UV-spectrophotometer. The temperature was scanned at a heating rate of 1°C/min at temperatures between 20°C and 90°C. Absorbance values were recorded with an average time of 0.5s and data were collected every 0.1°C. Samples contained 1μM pre-mRNA/gRNA hybrid RNA in 10mM Na-cacodylate pH6.8, 65mM NaCl. Half-maximal melting temperatures (T_m_) were calculated from 1^st^-derivative plots of absorbance versus temperature δA_260_/δT=f(T) (Katari et al., 2013).

### Ribonucleolytic degradation

The RNase-sensitivity of unmodified and phosphorothioate-modified 5’-Cl oligoribonucleotides (U-deletion: 5’-FAM_Cl22, 5’-FAM_synCl17, 5’-FAM_modCl17; U-insertion: 5’-TAMRA-Cl18, 5’-FAM_synCl14, 5’-FAM_modCl14) was tested by incubation of 0.4pmol RNA-oligonucleotide with 0.2pmol of editosomes in 30µL EB for 3h at 27°C. Samples were processed and analyzed as described above.

## Results and Discussion

### Starting analysis - Characterization of fluorophore-labeled pre-edited mRNA/gRNA hybrid RNAs

The established U-insertion/U-deletion *in vitro* RNA-editing assay (Seiwert and Stuart, 1994, Kable et al., 1996; Stuart et al., 1998) represents a chemically simplified experimental system in which both, the substrate pre-edited mRNAs as well as the *trans*-acting gRNAs are mimicked by short oligoribonucleotides (13-22nt) (Fig. 1). Furthermore, only single editing sites are embedded in each of the two pre-mRNAs, which in the case of the U-insertion assay is edited in a gRNA-dependent manner by incorporating 3 U-nucleotides (nt). During the U-deletion reaction 4 U-nt are removed. Furthermore, in order to sidestep the rate-limiting endonucleolytic cleavage of the pre-mRNA molecules, the corresponding oligoribonucleotides are “pre-cleaved”. This generates 5’- and 3’-pre-mRNA cleavage fragments, which are abbreviated as 5’-Cl and 3’-Cl (Fig. 1). The 5’-Cl oligoribonucleotides are typically 5’-(^32^P)-labeled to enable a radioactive readout of the assay. To convert the system into a fluorescence-based assay we covalently attached different fluorogenic side groups to either the 5’- or 3’-terminal phosphate groups of the two Cl-oligoribonucleotides. 5-Carboxytetramethylrhodamine (5-TAMRA, 430g/mol, λ_ex_ 546nm, λ_em_ 579nm), 6-carboxy-fluorescin (FAM, 750g/mol, λ_ex_ 492nm, λ_em_ 517nm) and 6-hexachlorofluorescein (HEX, 680g/mol, λ_ex_ 535nm, λ_em_ 556nm) were chosen as fluorophores. The coupling was performed post-synthetically using a 6-carbon atom spacer to minimize sterical clashes between the fluorophore moieties and the RNA-backbone. Since RNA-modifications can alter the physico-chemical properties of polynucleotides especially in the case of highly hydrophobic side-groups, we analyzed whether the different fluorophores affect the formation of the pre-mRNA/gRNA-hybrid RNAs. Antiparallel base-pairing between the pre-mRNA Cl-fragments and the corresponding gRNAs represents the first step in both *in vitro* editing reactions and the formation of the bi- and trimolecular RNA-hybrid molecules can be scrutinized by native polyacrylamide gel-electrophoresis. As shown in Fig. 2A, all non-modified oligoribonucleotides anneal rapidly *i.e*. within 10min and to completion (≥97%). No difference between the U-insertion and U-deletion RNA-hybrids is detected. The same result is obtained for all fluorophore-modified oligoribonucleotide complexes (Fig. 2B/C). Independent of the type of fluorophore (TAMRA, FAM, HEX) and independent of the covalent attachment site (5’ or 3’) all possible bi- and trimolecular pre-mRNA/gRNA complexes are formed with yields ≥95%. This verifies that both, the kinetic behaviour and the hydrogen bonding capacity of the different oligoribonucleotides are not affected by any of the fluorophore substituents. To broaden the analysis we further interrogated the thermodynamic stability of the fluorophore-modified hybrid RNAs. For that we recorded UV-melting profiles between 20°C and 90°C. Fig. 3 shows a representative result. As demonstrated before (Katari et al., 2013), non-modified versions of the insertion and deletion pre-mRNA/gRNA hybrids melt with two well separated transitions with T_m_-values of 68±1.5°C and 47±1.5°C for the U-insertion complex and 64±1.5°C and 55±1.5°C for the U-deletion hybrid. Importantly, all fluorophore-substituted versions of the two trimolecular RNA-complexes show qualitatively and quantitatively identical profiles. The derived T_m_-values deviate ≤1°C from the non-fluorophore substituted counterparts and as before, this behaviour is affected neither by the chemical nature nor by the structural positioning of the fluorophore moieties. For a complete list of all measured T_m_-values see supplementary Table S1.

**Figure 2.**
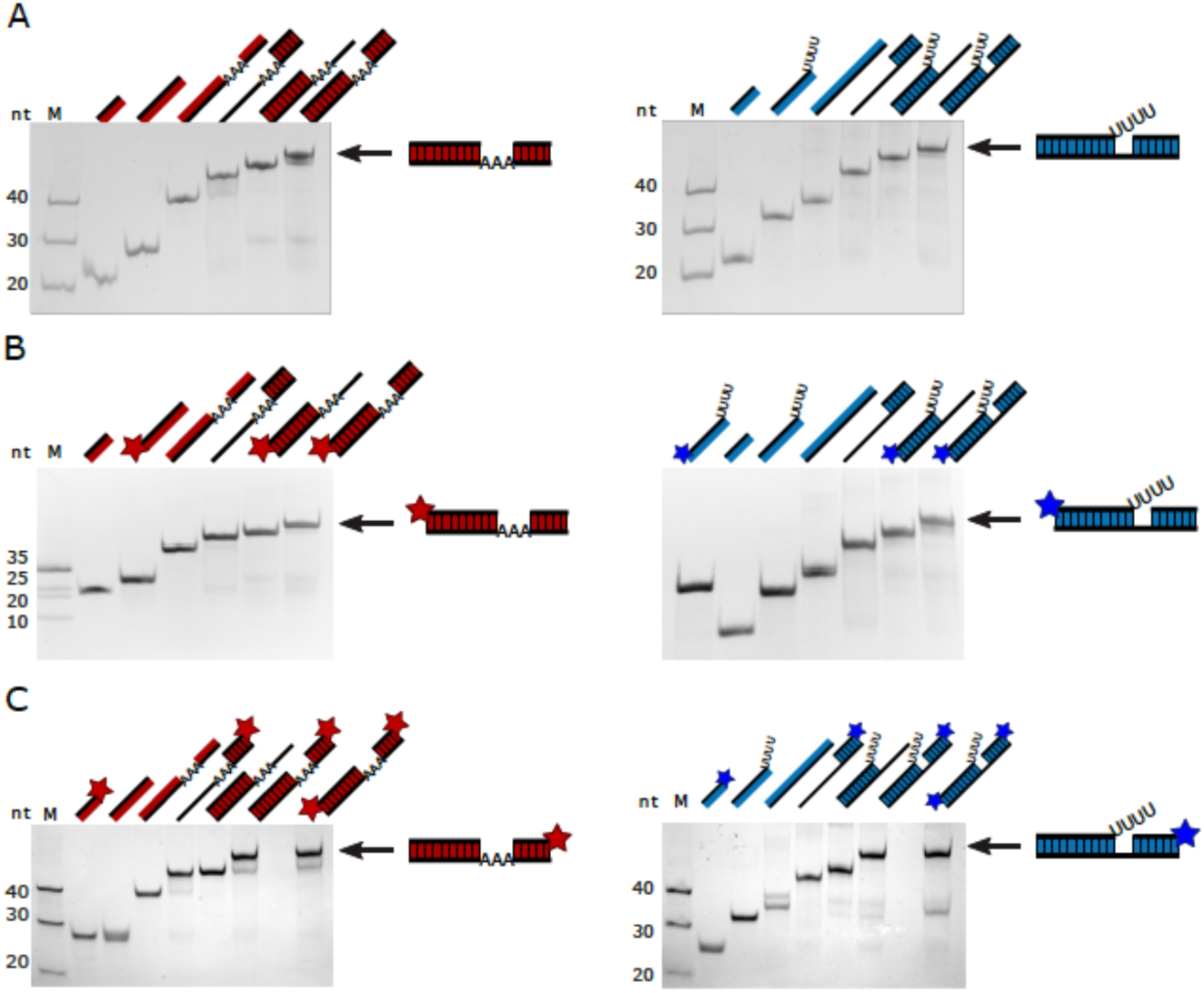
Pre-mRNA/gRNA-hybrid formation. Gel-electrophoretic analysis of the formation of trimolecular pre-mRNA/gRNA hybrid RNAs comparing non-modified RNA-oligonucleotides (A) with single 5’- or 3’-fluorophore-labeled RNAs (B, C) and dual-modified (5’ and 3’) RNAs (C). U-insertion RNAs are in red. U-deletion RNAs are in blue. All gel-electrophoretic separations show the 5’-pre-mRNA cleavage fragment (5’-Cl), the 3’-pre-mRNA cleavage fragment (3’-Cl) and the corresponding gRNA-oligoribonucleotide next to the two bimolecular complexes (5’-Cl/gRNA; 3’-Cl/gRNA) and the final trimolecular (5’-Cl/3’-Cl/gRNA) annealing product (arrow). M=marker, nt=nucleotides.

**Figure 3.**
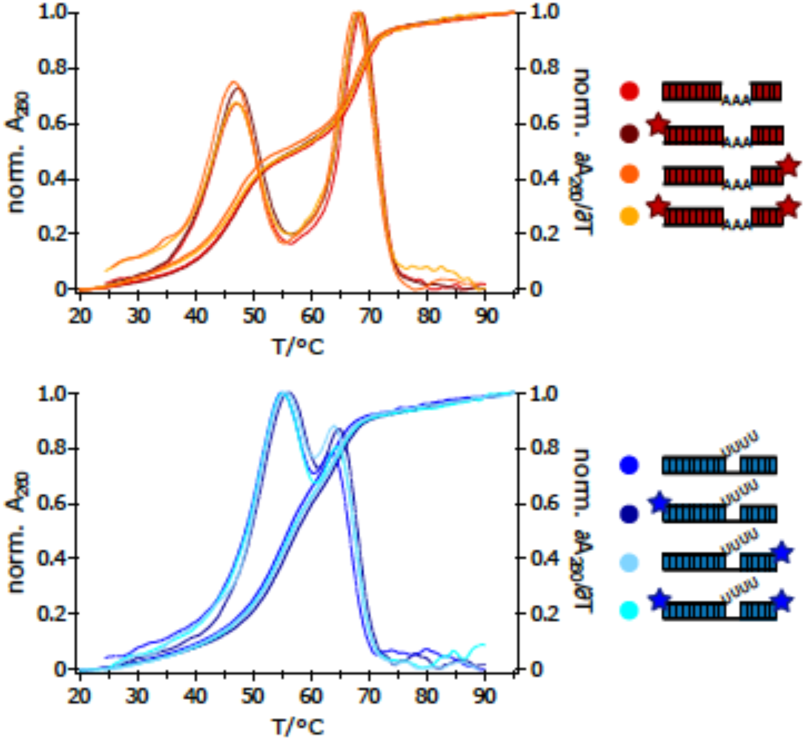
Thermodynamic stabilities of pre-mRNA/gRNA-hybrid RNAs. Comparison of the UV-melting (A_260_=f(T)) and 1^st^-derivative profiles (δA_260_/δT=f(T)) of non-labeled and fluorophore-labeled pre-mRNA/gRNA hybrid RNAs. Red: U-insertion pre-mRNA/gRNA hybrid molecules. Blue: U-deletion pre-mRNA/gRNA hybrid RNAs. Fluorophore-positions are annotated as stars.

### FIDE - Fluorescence-based Insertion/Deletion RNA-Editing

Next, we analyzed whether the fluorescently labeled pre-mRNA/gRNA hybrid RNAs are recognized by editosomes to act as *in vitro* RNA-editing substrates. For that we incubated fluorophore-modified insertion- and deletion-type pre-mRNA/gRNA hybrid RNAs with purified editosomes. Since the processing reaction is catalyzed in multiple enzyme-driven steps, we analyzed both, multiple turnover (editosomes<RNA-substrate) as well as single turnover conditions (editosomes≥RNA-substrate). This enabled us to identify the products of the catalytic conversion as well as all intermediates and side-products. After the reaction, samples were analyzed by automated capillary electrophoresis (CE) coupled to a laser-induced fluorescence (LIF) readout. Fig. 4A shows representative electropherograms in which the recorded fluorescence intensity (FI) in relative fluorescence units (RFU) is plotted as a function of the electrophoretic migration time (EM) in seconds. The U-deletion experiment was conducted with a FAM-derivatized pre-mRNA/gRNA hybrid and the electropherogram shows all 10 expected peaks: the 5’-Cl input RNA (+4U), all U-deletion intermediates of the 5’-Cl fragment as well as the fully edited and ligated -4U product. Additional RNA-species include the unedited 5’Cl/3’Cl-ligation product and all partially edited mRNAs in which only 1, 2 or 3 U’s have been deleted. In a similar fashion, the U-insertion assay was performed with a TAMRA-substituted pre-mRNA/gRNA hybrid. Fig. 4A shows all resolved RNA-species including the reaction educt, the +3U-editing product and the +1U, +2U reaction intermediates. Since the individual RNAs are resolved with nucleotide resolution, a relative quantification of every RNA-species was derived by peak integration. This enabled the calculation of percental RNA-editing activities (Fig. 4A) or of specific RNA-editing activities per mass of editosomes. An absolute (molar) quantification was accomplished by spiking the samples with known quantities (15fmol) of two 11nt and 44nt long calibration oligoribonucleotides (HEX_Low11, FAM_High44). Ultimately the data can be used to derive RNA-editing activity units (EU) defined as the catalytic conversion of 1pmol pre-mRNA/gRNA-hybrid RNA into fully edited mRNA in 1 hour at 27°C. To test the reproducibility of the fluorescence-based assay, we compared up to nine technical and six biological replicates including editosome preparations from different *T. brucei* genetic backgrounds. The data were analyzed in a bivariate correlation analysis to calculate Pearson’s correlation coefficients (*p*). Fig. 4B shows a summary of the results. Technical replicates are characterized by *p*-values ≥0.99 and biological replicates by *p*-values ≥0.89 demonstrating a high level of experimental robustness. A comparison of the fluorophore CE/LIF-based *in vitro* assays with the standard radioactive, slab gel-based assays resulted in *p*-values ≥0.86 (suppl. Fig. S2).

**Figure 4.**
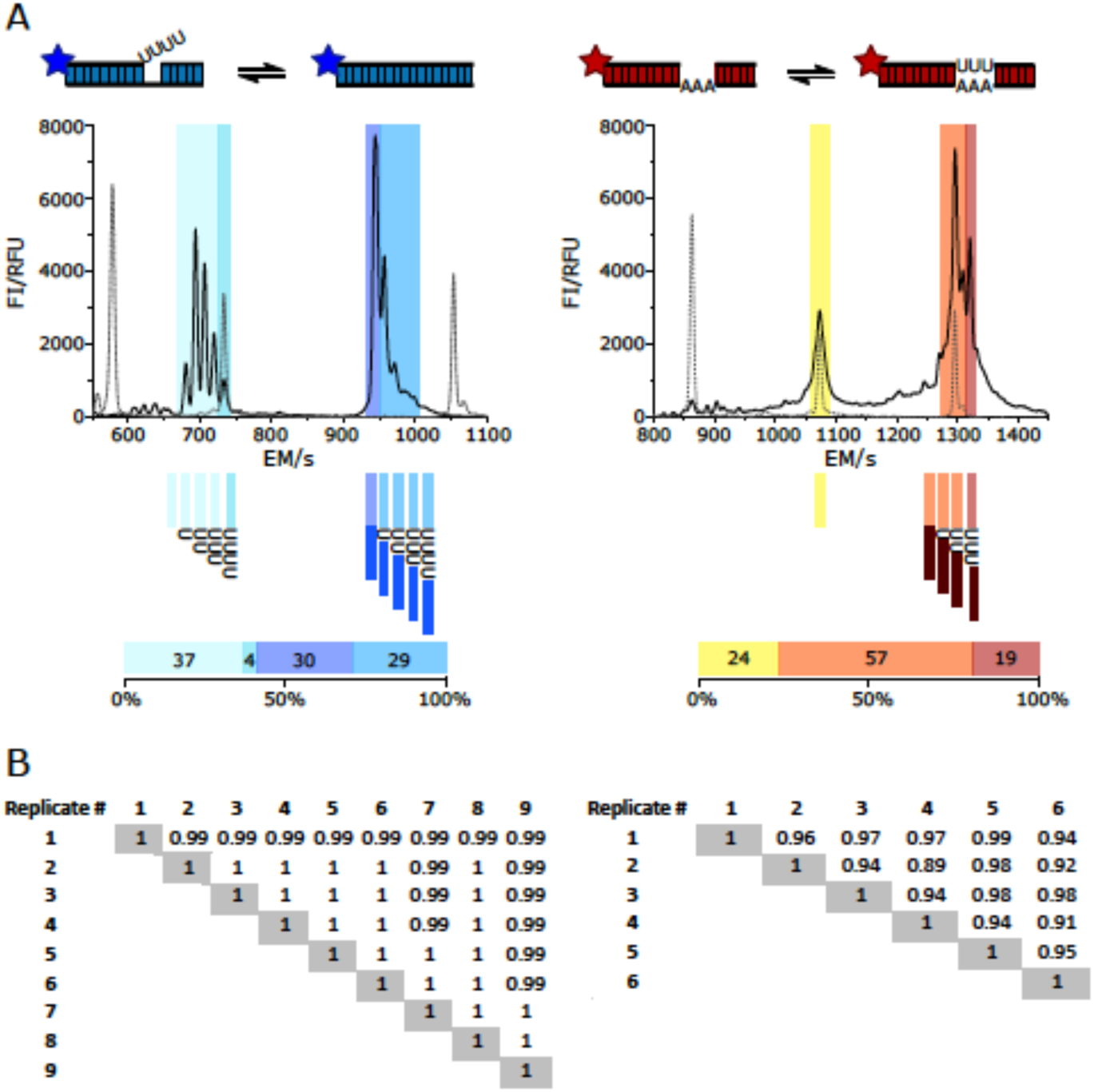
Fluorescence-based insertion/deletion RNA-editing (FIDE). (A) Left: capillary electrophoretic (CE)-separation of the products of a U-deletion *in vitro* RNA-editing reaction using a 5’-FAM fluorophore-modified pre-mRNA cleavage fragment (5’-FAM_Cl22). The different shadings in blue highlight the input 5’-FAM_Cl22-fragment, all reaction intermediates as well as the fully edited -4U reaction product. Right: CE-trace of a U-insertion editing reaction using a 5’-TAMRA-fluorophore-modified pre-mRNA cleavage fragment (5’-TAMRA_Cl18). Shadings in yellow/red mark the input 5’-TAMRA_Cl18 pre-mRNA fragment, reaction intermediates and the fully edited +3U insertion product. Peak area integration allows the calculation of percental quantities for every intermediate and product as shown in the bar-plots below the electropherograms. FI=fluorescence intensity, RFU=relative fluorescence units; EM=electrophoretic migration time in sec. Dashed lines/peaks (from left to right) represent the electrophoretic elution positions of the 11nt marker-oligoribonucleotide HEX_Low11, the unprocessed 5’-Cl oligoribonucleotides and the 44nt marker-oligoribonucleotide FAM_High44. (B) Left: Pairwise comparison of Pearson correlation coefficients (*p*) of 9 technical FIDE-assay replicates. Right: Pairwise comparison of *p*-values from 6 biological FIDE-assay replicates.

### Optimization I - multiplexing and downscaling

The structure-conservative characteristics of the fluorophore-substituted oligoribonucleotides encouraged us to investigate whether the assay system can be improved by multiplexing *i.e*. by using multiple fluorophore modifications. Since modern CE/LIF-instruments can monitor up to five different fluorophores in one electrophoresis run we specifically addressed two scenarios: 1. a dual fluorophore-labeling approach of the two pre-mRNA-mimicking oligoribonucleotides and 2. a side-by-side analysis of both, U-insertion and U-deletion editing in one reaction vial. Representative examples of the generated data-sets are shown in Fig. 5. By using a 5’-FAM-substituted U-deletion 5’-Cl fragment in combination with a 3’-TAMRA-modified U-deletion 3’-Cl oligoribonucleotide the conversion of both pre-mRNA fragments into the same partially and fully-edited reaction products is unequivocally demonstrated (Fig. 5A). Both, the FAM- and TAMRA-signals overlap for all partially and fully edited U-deletion products, while the two educts and all intermediates are well separated from each other. At the same time, the experiment scrutinizes the faith of both pre-mRNA fragments and corroborates that only the 5’-pre-mRNA fragment acts as source for the -1U, -2U, -3U and -4U intermediates. Similarly, by using a 5’-FAM-labeled U-deletion and a 5’-TAMRA-substituted U-insertion pre-mRNA/gRNA hybrid we demonstrate that both editing reactions can be performed in “one-pot” (Fig. 5B). The two CE-tracings are not influenced by each other and no qualitative or quantitative difference in comparison to the separate CE-runs is observed.

**Figure 5.**
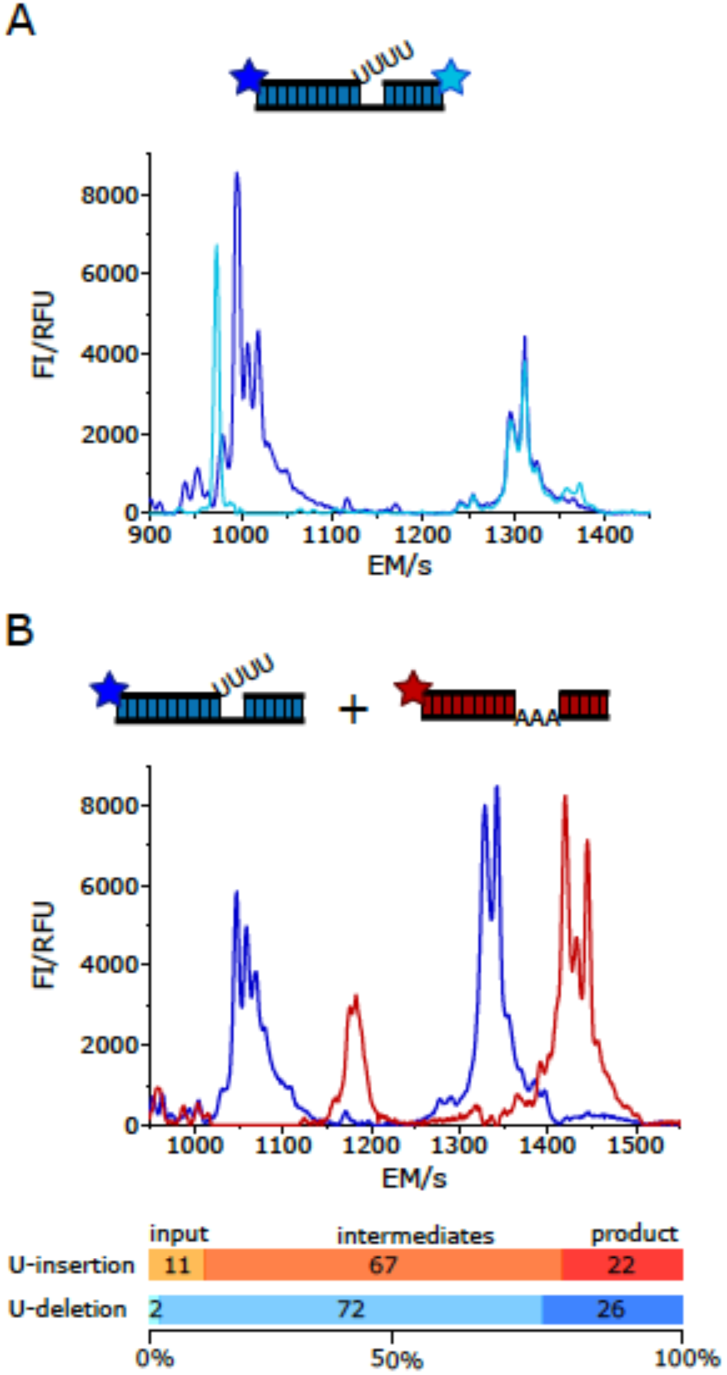
FIDE-multiplexing. (A) Cartoon of a dual fluorophore-labeled U-deletion RNA-editing substrate RNA, assembled by using a 5’-FAM-labeled 5’-pre-mRNA Cl-fragment (5’-FAM_Cl22) and a 3’-TAMRA-modified 3’ Cl-fragment (3’-Cl15_TAMRA). The electropherogram shows the FAM-derived CE/LIF-trace in dark blue and the TAMRA-based trace in light blue. The data demonstrate that the reaction pathway of the two pre-mRNA fragments can be investigated in one substrate molecule however, independent from each other. Other multiple fluorophore-labeling strategies are also possible. The two CE/LIF-traces further show that both fluorescence signals converge for the product peak and all ligated intermediates confirming the re-ligation of the 5’- and 3’-pre-mRNA Cl-fragments after the catalytic conversion. (B) “One pot” U-insertion and U-deletion RNA-editing. The cartoon depicts a trimeric 5’-FAM-modified U-deletion RNA-substrate in blue and a trimeric 5’-TAMRA-substituted U-insertion RNA-substrate in red. Both hybrid-RNAs were edited in the same reaction vial followed by automated CE/LIF-detection. Bar-plots show the percental abundance of all reaction intermediates and products in relation to the amount of input-RNA. The ability to monitor both editing reactions in one sample further reduces the sample size in HTS-screening experiments. FI=fluorescence intensity, RFU=relative fluorescence unit, EM=electrophoretic migration time in sec.

As a follow up we tested whether the two *in vitro* reactions can be downscaled to tailor the assay for high-throughput screening applications. This was done in a stepwise fashion using the signal to noise ratio (S/N) of the CE-electropherograms as a proxi. As lower limit we determined 2.5-10fmol of each, editosomes and fluorophore-modified pre-mRNA/gRNA-hybrid in a reaction volume of 4µL. At these conditions signal intensities remained about 300-fold over background for high intensity peaks and 50-fold over background for low intensity peaks. Because of the minute amounts of protein and RNA in the samples, reactions were stopped by directly adding formamide thereby eliminating the time-consuming phenol and EtOH-treatment steps. Furthermore, the small assay volume enables the usage of multiwell plates, which makes the test suitable for HTS-experiments. As a statistical HTS-quality score (Zhang et al., 1999) we determined a mean screening window coefficient (Z’) of 0.6 confirming the aptness of the assay for HTS-screening purposes. Using the downscaled assay conditions, it is possible to quantitatively monitor 5000 chemical compounds on their influence on both RNA-editing reactions with only 10µg (12.5pmol) of editosomes and 2µg (2×50pmol) of fluorophore-modified pre-mRNA/gRNA-hybrid RNAs in just 78h, using a 96-capillary CE-instrument.

Lastly, we scrutinized the kinetic of the catalytic conversion. Supplementary Fig. S3 shows as an example the CE/LIF-traces of a U-deletion editing reaction, incubated for 5min up to 180min. Reaction intermediates and products can be identified as early as 5min. At 50min, the processed/input RNA-ratio approaches a value of 50% demonstrating that the reaction time can be shortened much below the standard 3hours of incubation.

### Optimization II - RNA-substrate chemistry

To further improve the FIDE-assay we focused on two additional aspects: 1. the ribonucleolytic stability of the substrate RNAs and 2. possible sequence context effects, especially if both editing reactions are executed side-by-side in the same reaction vial. Since editosomes are purified from whole cell lysates co-purifying and/or contaminating RNases can be challenging (for an example see suppl. Fig S4A). The problem is even more pronounced if only partially purified mitochondrial (mt)-extracts are used as editosome source, despite the fact that mt-extracts are better suited for large-scale screening purposes due to time and budgetary concerns. To reduce the RNase sensitivity and, at the same time, to avoid RNA-sequence biases we designed a non-natural pre-mRNA/gRNA-hybrid RNA that includes multiple, RNase-resistant phosphorothioate bonds. The trimeric RNA-molecule is of identical sequence for both, the U-insertion and U-deletion reaction and differs only at the editing site where either 3 U-nt are inserted or deleted (Fig. 6A).

**Figure 6.**
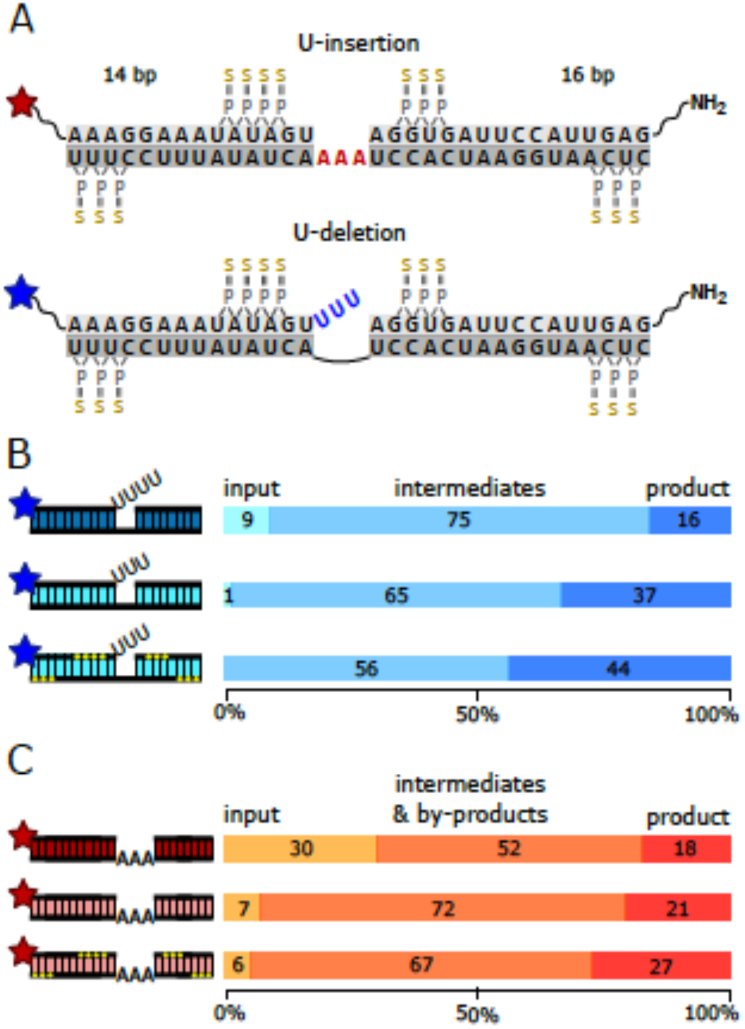
Pre-mRNA/gRNA hybrid-RNA optimization. (A) Sequence representation of the non-natural pre-mRNA/gRNA hybrid RNAs. Top: U-insertion hybrid-RNA. Bottom: U-deletion hybrid-RNA. The 5’ and 3’ pre-mRNA cleavage fragments are in light grey, gRNA-sequences in dark grey. The 14bp and 16bp-long helical RNA-elements flanking the U-insertion and U-deletion editing sites are identical in both trimeric RNAs. Guide RNA nucleotides dictating the insertion of 3 U’s are in red. The 3 U’s that are deleted are in blue. Phosphorothioate modifications are highlighted in yellow and stars show fluorophore attachment sites. (B) Cartoons of U-deletion-type pre-mRNA/gRNA hybrid RNAs. Top: standard hybrid-RNA (dark blue); center: non-natural hybrid-RNA (light blue); bottom: non-natural, phosphorothioate (PS)-modified RNA (light blue, yellow dots indicate PS-positions). Bar-plots to the right summarize the FIDE-assay results specifying the percental amounts of U-deletion product, editing intermediates and input-RNA. (C) Cartoons of U-insertion-type pre-mRNA/gRNA hybrid-RNAs. Top: standard hybrid-RNA (dark red); center: non-natural RNA-hybrid (light red); bottom: non-natural, phosphorothioate (PS)-modified RNA (light red, yellow dots indicate PS-positions). Bar-plots to the right summarize the FIDE-assay results specifying the percental amounts of U-insertion product, editing intermediates and input-RNA. For both, the U-deletion and the U-insertion reaction, the non-natural, PS-modified RNAs represent the most efficient substrate molecules. Stars denote fluorophore positions.

The synthetic RNA-sequence was selected from a randomized sequence pool and has no match in the mitochondrial genome of African trypanosomes. Phosphorothioates (PS) substitute a sulfur atom for a non-bridging oxygen in the phosphate backbone and this renders the internucleotide linkage nuclease resistant. Altogether 13 phosphorothioates were co-synthetically introduced into the trimolecular hybrid-RNAs: 4, respectively 3 PS were positioned on either side of the pre-mRNA editing domains and 6 additional phosphorothioates were placed next to the two termini of the gRNA-oligonucleotides (Fig. 6A). Incubation of the PS-derivatized oligoribonucleotides with editosomes in the absence of gRNAs verified their RNase-resistant characteristics (suppl. Fig. S4B/C). While up to 70% of the unmodified 5’-pre-mRNA oligonucleotides were degraded within 3 hours at 27°C, 70-80% of the PS-modified RNAs remain stable over the same period. Furthermore, we verified that the synthetic, PS-modified oligoribonucleotides are able to form trimeric pre-mRNA/gRNA complexes with reaction kinetics and thermodynamic stabilities that match their non-modified counterparts (suppl. Fig. S5, S6 and Table S1). And finally, we tested the PS-substituted, non-natural pre-mRNA/gRNA hybrids as editing substrates. Fig. 6B/C demonstrate that both RNA-hybrids are properly processed by editosomes generating all expected fully and partially edited products and intermediates. A comparison of the product/input (p/i)-ratio identifies that the non-natural pre-mRNA/gRNA-hybrids outperform the trypanosome-based pre-mRNA/gRNA sequences between 10 to 20-fold, which together with their improved RNase-stability makes them well suited as substrates in HTS-experiments.

### Validation - screening for inhibitory UTP-analogs

To demonstrate the high-throughput potential of the FIDE-assay, we performed a small-scale, proof-of-principle screening experiment. For that we made use of the non-natural U-insertion pre-mRNA/gRNA-substrate described above and examined whether U-nucleotide analogs might interfere with the U-insertion reaction. Both, pyrimidine ring-modified compounds (5-CH_3_-UTP, 5-Br-UTP, 5-OH-UTP, 4-S-UTP, 2-S-UTP, 5-biotin-UTP, 5-CH_3_N_3_-UTP etc.) as well as ribose-modified analogs such as 2’-CH_3_O-UTP, 2’-NH_2_-UTP, 2’-F-UTP, 2’-N_3_-UTP and Ara-UTP were tested. A list of all compounds is shown in Fig. 7A.

**Figure 7.**
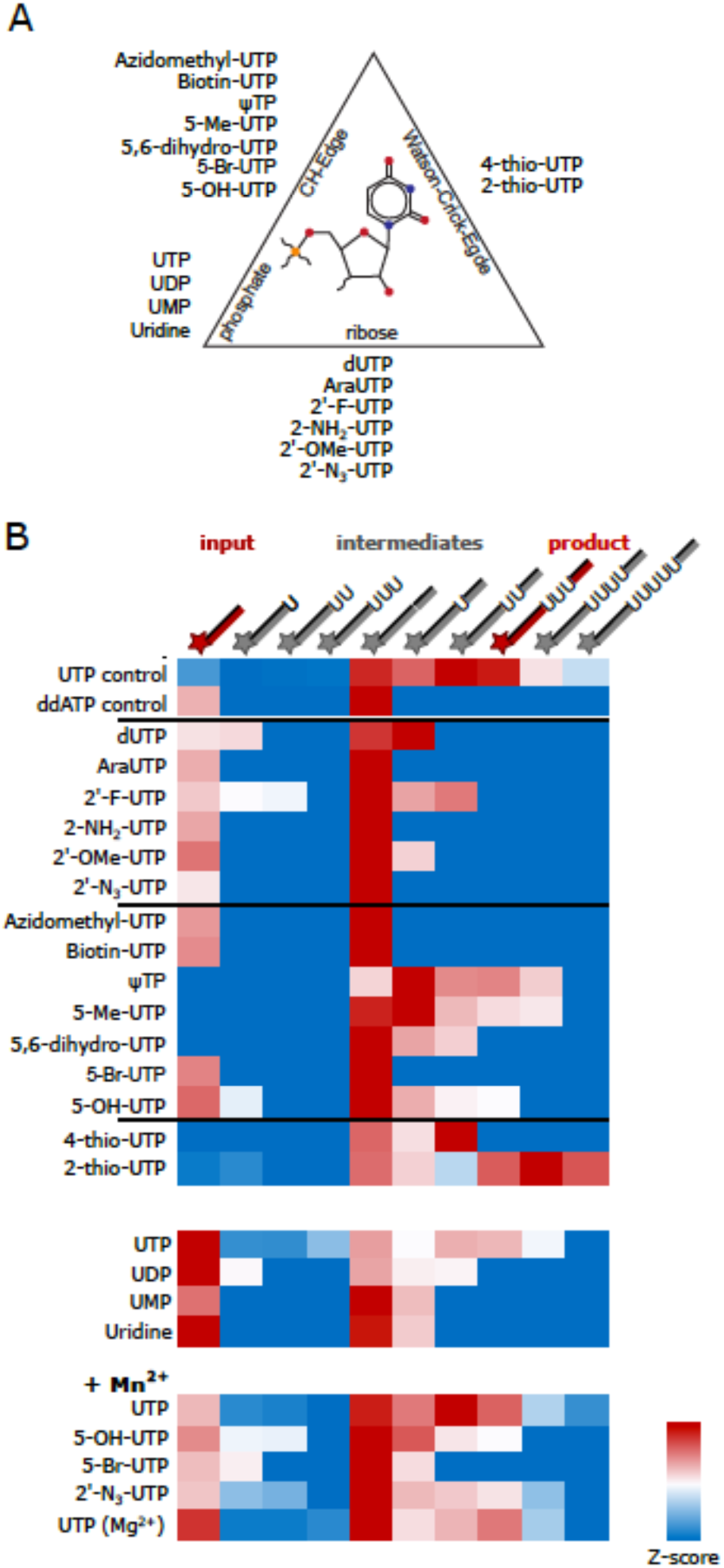
Inhibition of U-insertion RNA-editing by UTP-analogs. (A) Tested UTP-analogs are listed next to a ball-and-stick representation of UMP to positionally categorize the different compounds as CH edge-, Watson/Crick edge-, phosphate- or ribose-interfering. (B) Heat-map (Z-score) representation of the FIDE-assay results ranging from blue (low abundance) to red (high abundance) for the reaction input, all reaction intermediates and the U-insertion product. The heat-map block at the bottom summarizes the incorporation of UTP-analogs in the presence of Mn^2+^ instead of Mg^2+^.

In addition, we probed the influence of Mg^2+^- and Mn^2+^-cations in the reaction. Altogether 26 samples were tested, which on a 48 multicapillary CE/LIF-instrument can be assayed in just 1.5h generating 260 quantitative data points on the formation of the fully edited product and on every reaction intermediate. As shown in Fig. 7B the data confirm that UTP is the preferred substrate of the reaction. Furthermore, the data reveal that the ribose 2’-OH group is more important in the catalytic conversion than the majority of substituents at the CH- and Watson/Crick-edges of the U-nucleobase. All 2’-ribose modifications inhibit the formation of the fully edited (+3U) product and only 2’-F-UTP is to some degree tolerated (forming a +2U product). By contrast, multiple responses were detected for all base-modifications. They range from complete inhibition by 5-Br-UTP to different degrees of misediting by 5-CH_3_-UTP (+4Us) and 2-S-UTP (+4 and +5U’s). Mn^2+^-ions have an enhancing effect, which can result in the incorporation of otherwise non-inserted UTP-analogs. Together, the data demonstrate the versatility of the FIDE-*in vitro* system by enabling a comprehensive and quantitative inhibitor screening in a very short time.

## Conclusion

In summary, we have converted the standard *in vitro* assay to monitor editosome function in African trypanosomes and related organisms into a high-throughput analysis format. Core attribute of the revised assay is the usage of fluorophore-labeled RNA-editing substrate RNAs. This enables the automated electrophoretic separation of the products of the catalytic conversion using state-of-the-art, high-throughput capillary electrophoresis instruments coupled with laser-induced fluorescence readout systems. As such the assay generates quantitative data of all educts, intermediates and products and by using multicapillary instruments it can be performed in a highly parallel format. We optimized the assay by downscaling the required material quantities as well as the reaction volume to adapt the procedure to all available multiwell-plate formats. Additional improvements include the usage of non-natural, RNase-resistant RNA-substrates, which enable large-scale screening experiments using crude mitochondrial extracts instead of highly purified editosome preparations. We also verified that the experimental setup tolerates multiplex-type, fluorophore-labeling strategies, which expands the parallel analysis-capacity of the assay further. As demonstrated in a pilot screening experiment, the assay is applicable to larger sample-size studies to probe specific mechanistic aspects of the RNA-editing reaction. However, the assay is especially well suited to conduct large-scale, high-throughput screening experiments to identify small molecule inhibitors. Since RNA-editing is an essential biochemical pathway in African trypanosomes, an assay format that permits a robust, quantitative as well as time- and material-resourceful analysis should be helpful in the search for novel therapeutics to challenge African trypanosomiasis and related diseases.

## Supplementary Data

Supplementary data are available online.

## Acknowledgements

We thank Reza Salavati, Inna Aphasizheva and Ruslan Aphasizhev for transgenic trypanosome strains. Robert Knieß and Andreas Völker are acknowledged for experimental help and Elisabeth Kruse for discussions.

## References

Aphasizheva I, Alfonzo J, Carnes J, Cestari I, Cruz-Reyes J, Göringer HU, Hajduk S, Lukeš J, Madison-Antenucci S, Maslov DA, McDermott SM, Ochsenreiter T, Read LK, Salavati R, Schnaufer A, Schneider A, Simpson L, Stuart K, Yurchenko V, Zhou ZH, Zíková A, Zhang L, Zimmer S, Aphasizhev R. 2020. Lexis and grammar of mitochondrial RNA processing in trypanosomes. Trends Parasitol. 36, 337–355.

Baker N, de Koning HP, Mäser P, Horn D. 2013. Drug resistance in African trypanosomiasis: the melarsoprol and pentamidine story. Trends Parasitol. 29:110–118.

Barrett MP. 2018. The elimination of human African trypanosomiasis is in sight: Report from the third WHO stakeholders meeting on elimination of Gambiense human African trypanosomiasis. PLoS Negl Trop Dis. 12, e0006925.

Barrett MP, Vincent IM, Burchmore RJ, Kazibwe AJ, Matovu E. 2011. Drug resistance in human African trypanosomiasis. Future Microbiol. 6, 1037–1047.

Benne R, Van Den Burg J, Brakenhoff JPJ, Sloof P, Van Boom JH, Tromp MC. 1986. Major transcript of the frameshifted coxll gene from trypanosome mitochondria contains four nucleotides that are not encoded in the DNA. Cell 46, 819–826.

Blum B, Bakalara N, Simpson L. 1990. A model for RNA editing in kinetoplastid mitochondria: RNA molecules transcribed from maxicircle DNA provide the edited information. Cell 60, 189–198.

Böhm C, Katari VS, Brecht M, Göringer HU. 2012. *Trypanosoma brucei* 20S editosomes have one RNA substrate-binding site and execute RNA unwinding activity. J Biol Chem. 287, 26268–26277.

Brun R, Schönenberger M. 1979. Cultivation and *in vitro* cloning or procyclic culture forms of *Trypanosoma brucei* in a semi-defined medium. Acta Trop. 36, 289–292.

Brun R, Blum J, Chappuis F, Burri C. (2010) Human African trypanosomiasis. Lancet 375, 148–159.

Büscher P, Cecchi G, Jamonneau V. and Priotto G. (2017) Human African trypanosomiasis. Lancet 390, 2397–2409.

Burri C. 2020. Sleeping sickness at the crossroads. Trop Med Infect Dis. 5, 57.

Ernst NL, Panicucci B, Igo RP Jr, Panigrahi AK, Salavati R, Stuart K. 2003. TbMP57 is a 3’ terminal uridylyl transferase (TUTase) of the *Trypanosoma brucei* editosome. Mol Cell 11, 1525–1536.

Fairlamb AH, Horn D. (2018) Melarsoprol resistance in African trypanosomiasis. Trends Parasitol. 34, 481–492.

Field MC, Horn D, Fairlamb AH, Ferguson MAJ, Gray DW, Read KD, De Rycker M, Torrie LS, Wyatt PG, Wyllie S, Gilbert IH. 2017. Anti-trypanosomatid drug discovery: an ongoing challenge and a continuing need. Nat Rev Microbiol. 15, 447.

Gerace E, Moazed D. 2015. Affinity purification of protein complexes using TAP tags. Methods Enzymol. 559, 37–52.

Gilbert IH. 2014. Target-based drug discovery for human African trypanosomiasis: Selection of molecular target and chemical matter. Parasitology 141, 28–36.

Golas MM, Böhm C, Sander B, Effenberger K, Brecht M, Stark H, Göringer HU. 2009. Snapshots of the RNA editing machine in trypanosomes captured at different assembly stages *in vivo*. EMBO J. 28, 766–778.

Göringer, H.U. (2012) ‘Gestalt,’ composition and function of the *Trypanosoma brucei* editosome. Annu Rev Microbiol. 66, 65–82.

Hermann T, Schmid B, Heumann H, Göringer HU. 1997. A three-dimensional working model for a guide RNA from *Trypanosoma brucei*. Nucleic Acids Res. 25, 2311–2318.

Jamonneau V, Ilboudo H, Kaboré J, Kaba D, Koffi M, Solano P, Garcia A, Courtin D, Laveissière C, Lingue K, Büscher P, Bucheton B. 2012. Untreated human infections by *Trypanosoma brucei gambiense* are not 100% fatal. PLoS Negl Trop Dis. 6, e1691.

Kable ML, Seiwert SD, Heidmann S, Stuart K. 1996. RNA editing: a mechanism for gRNA-specified uridylate insertion into precursor mRNA. Science 273, 1189–1195.

Kala S, Moshiri H, Mehta V, Yip CW, Salavati R. 2012. The oligonucleotide binding (OB)-fold domain of KREPA4 is essential for stable incorporation into editosomes. PLoS One 7:e46864.

Katari VS, van Esdonk L, Göringer HU. 2013. Molecular crowding inhibits U-insertion/deletion RNA editing *in vitro*: consequences for the *in vivo* reaction. PLoS One 8, e83796.

Kennedy PGE. 2019. Update on human African trypanosomiasis (sleeping sickness). J. Neurol. 266, 2334–2337.

Leeder WM, Reuss AJ, Brecht M, Kratz K, Wachtveitl J, Göringer HU. 2015. Charge reduction and thermodynamic stabilization of substrate RNAs inhibit RNA editing. PLoS One 10, 1–17.

Liang S, Connell GJ. 2009. An electrochemiluminescent aptamer switch for a high-throughput assay of an RNA editing reaction. RNA 15, 1929–1938.

Liang S, Connell GJ. 2010. Identification of specific inhibitors for a trypanosomatid RNA editing reaction. RNA 16, 2435–2441.

Moshiri H, Salavati R. 2010. A fluorescence-based reporter substrate for monitoring RNA editing in trypanosomatid pathogens. Nucleic Acids Res. 38, 1–13.

Pollard VW, Harris ME, Hajduk SL. 1992. Native mRNA editing complexes from *Trypanosoma brucei* mitochondria. EMBO J. 11, 4429–4438.

Rigaut G, Shevchenko A, Rutz B, Wilm M, Mann M, Seraphin B. 1999. A generic protein purification method for protein complex characterization and proteome exploration. Nat. Biotechnol. 17, 1030–1032.

Ringpis GE, Aphasizheva I, Wang X, Huang L, Lathrop RH, Hatfield GW, Aphasizhev R. 2010. Mechanism of U insertion RNA editing in trypanosome mitochondria: the bimodal TUTase activity of the core complex. J Mol Biol. 399, 680–695.

Salavati R, Ernst NL, O’Rear J, Gilliam T, Tarun S, Stuart K. 2006. KREPA4, an RNA binding protein essential for editosome integrity and survival of *Trypanosoma brucei*. RNA 12, 819–831.

Salavati R, Moshiri H, Kala S, Shateri Najafabadi H. 2012. Inhibitors of RNA editing as potential chemotherapeutics against trypanosomatid pathogens. Int J Parasitol Drugs Drug Resist. 2, 36–46.

Seiwert SD, Stuart K. 1994. RNA editing: Transfer of genetic information from gRNA to precursor mRNA *in vitro*. Science 266, 114–117.

Stuart K, Kable ML, Allen TE,Lawson S. 1998. Investigating the Mechanism and Machinery of RNA Editing. Methods, 15, 3–14.

Vasa SM, Guex N, Wilkinson KA, Weeks KM, Giddings MC. 2008. ShapeFinder: A software system for high-throughput quantitative analysis of nucleic acid reactivity information resolved by capillary electrophoresis. RNA, 14, 1979–1990.

Zhang JH, Chung TD, Oldenburg KR. 1999. A simple statistical parameter for use in evaluation and validation of high throughput screening assays. J Biomol Screen. 4, 67–73.

Zimmermann S, Hall L, Riley S, Sørensen J, Amaro RE, Schnaufer A. 2016. A novel high-throughput activity assay for the *Trypanosoma brucei* editosome enzyme REL1 and other RNA ligases. Nucleic Acids Res. 44, 1–12.

