## Supplementary material for "Analyzing editosome function in high-throughput"

Cristian Del Campo, Wolf-Matthias Leeder, Paul Reißig, H. Ulrich Göringer
Molecular Genetics, Technical University Darmstadt, Schnittspahnstr. 10, 64287 Darmstadt, Germany

Supplementary Figures:

| | |
|---|---|
| Figure S1. | Oligoribonucleotide quality assessment |
| Figure S2. | Comparison of the ($^{32}$P)-isotope labeling-based *in vitro* RNA-editing assays to FIDE |
| Figure S3. | Kinetic of the catalytic conversion |
| Figure S4. | RNase stability |
| Figure S5. | Annealing of non-natural pre-mRNA/gRNA-hybrid RNAs. |
| Figure S6. | Thermodynamic stabilities of non-natural pre-mRNA/gRNA-hybrid RNAs. |

Supplementary Tables:

| | |
|---|---|
| Table S1. | Summary of half-maximal melting transitions ($T_m$-values) for the different pre-mRNA/gRNA-hybrid RNAs |

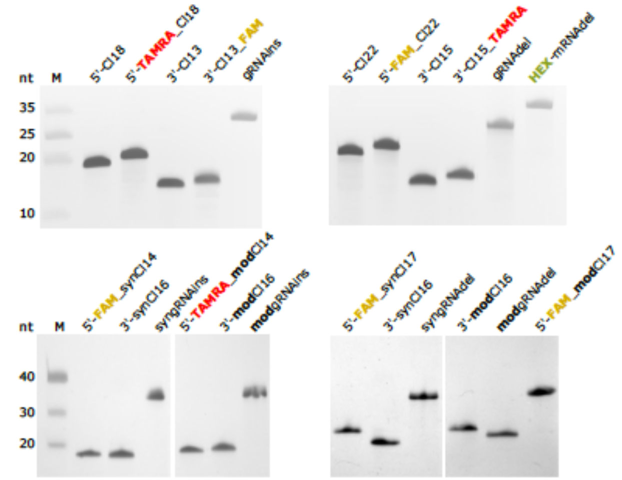


**Figure S1. Oligoribonucleotide quality assessment**. Chemically synthesized oligoribonucleotides were postsynthetically scrutinized by gel-electrophoresis in 8M urea-containing, 15% (w/v) polyacrylamide gels followed by Toluidine Blue O staining. Abbreviations and sequences of the different RNA-molecules are specified in the Materials and Method section. Fluorophore modifications (TAMRA, FAM, HEX) are coloured in red, yellow and green and phosphorothioate-modifictions (mod) are in bold. All RNA-preparations are ≥97% pure.

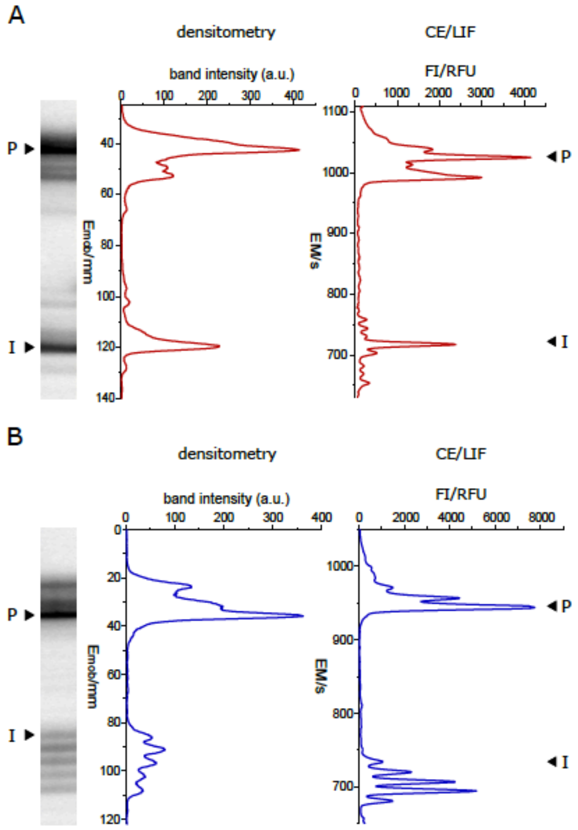


**Figure S2. Comparison of the isotope labeling-based *in vitro* RNA-editing assay to FIDE.** (A) U-insertion editing. (B) U-deletion editing. ($^{32}$P)-labeled RNA-reaction products were electrophoretically separated in 8M urea-containing 18% (w/v) polyacrylamide gels and were densitometrically quantified. Fluorophore-labeled substrate RNAs were analyzed by CE/LIF. (I)=input RNA. (P)=fully edited RNA-product. The experiments compare with Pearson correlation coefficients of $p$=0.87 for the U-insertion assays and with $p$=0.86 for the U-deletion assays. FI=fluorescence intensity, RFU=relative fluorescence unit, EM=electrophoretic migration time in seconds and $E_{mob}$=electrophoretic mobility in mm.

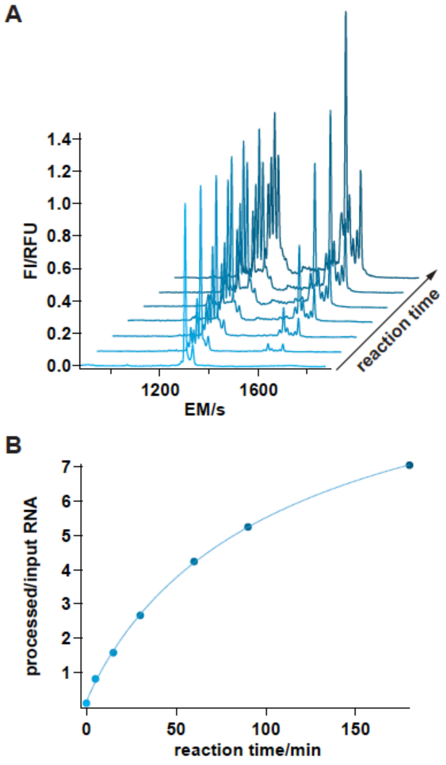


**Figure S3. Kinetic of the catalytic conversion.** Time course of a U-deletion FIDE-assay for reaction times of 0, 5, 15, 30, 60, 90 and 180min. (A) CE/LIF-traces of the different samples from 0min (front) to 180min (back). EM=electrophoretic migration time in seconds**.** FI=fluorescence intensity. RFU=relative fluorescence unit. (B) Plot of the processed/input RNA-ratio over time. Data points were curve-fitted to the Hill-equation with a chi-square of 0.015.

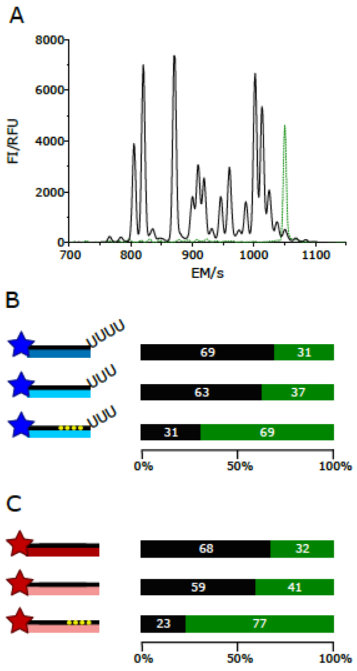


**Figure S4. RNase stability.** Ribonucleolytic degradation of fluorophore-modified 5'-pre-mRNA cleavage fragments (Cl) upon incubation with editosomes. (A) CE-trace (black) of a fully degraded 5'-Cl fragment (dashed trace in green: input 5'-Cl-RNA). (B) U-deletion 5' Cl-fragments (cartoons in blue): 5'-FAM_Cl22, 5'-FAM_synCl17, 5'-FAM_modCl17. (C) U-insertion 5' Cl-fragments (cartoons in red): 5'-TAMRA-Cl18, 5'-FAM_synCl14, 5'-FAM_modCl14. Top: standard 5'-Cl RNA-oligonucleotides. Centre: non-natural 5'-Cl RNA-oligonucleotides. Bottom: non-natural, phosphorothioate (PS)-modified 5'-Cl RNA-oligonucleotides. Yellow dots indicate PS-positions. Bar-plots display the percentage of intact and degraded RNA after a 3h incubation at 27°C.

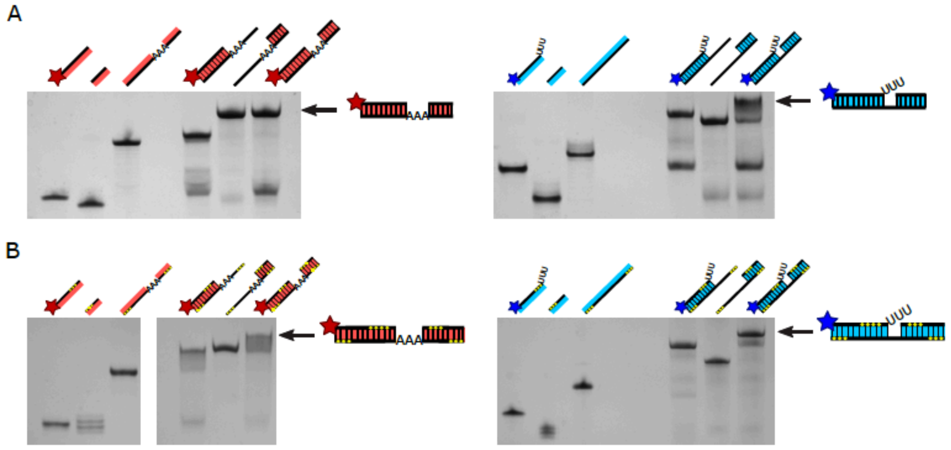


**Figure S5. Annealing of non-natural pre-mRNA/gRNA-hybrid RNAs.** Gel-electrophoretic analysis of the formation of trimolecular pre-mRNA/gRNA hybrid RNAs for the non-natural pre-mRNA/gRNA editing substrate (A) and the non-natural, phosphorothioate (PT)-modified editing substrates (B). U-insertion RNAs are in red, U-deletion RNAs in blue. Stars show the positions of the fluorophore subtituents and yellow dots PS-positions. All gel-electrophoretic separations show the 5'-pre-mRNA cleavage fragment (5'-Cl), the 3'-pre-mRNA cleavage fragment (3'-Cl) and the corresponding gRNA-oligoribonucleotide next to the two bimolecular complexes (5'-Cl/gRNA; 3'-Cl/gRNA) and the final trimolecular (5'-Cl/3'-Cl/gRNA) annealing product (arrow).

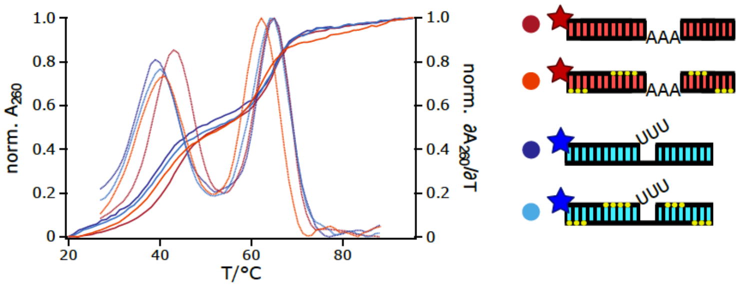


**Figure S6. Thermodynamic stabilities of non-natural pre-mRNA/gRNA-hybrid RNAs.** Comparison of the UV-melting ($A_{260}$=f(T)) and $1^{st}$-derivative profiles ($\delta A_{260}/\delta T$=f(T)) of non-natural pre-mRNA/gRNA hybrid-RNAs and non-natural, phosphorothioate (PT)-modified pre-mRNA/gRNA hybrid-RNAs. Red/orange: U-insertion RNAs. Dark blue/light blue: U-deletion RNAs. Yellow dots mark PT-modifications. Fluorophore positions are shown as stars. Halfmaximal melting temperatures ($T_m$) are listed in supplementary Table S1.

**Table S1. Summary of half-maximal melting transitions ($T_m$-values) for the different pre-mRNA/gRNA-hybrid RNAs.** Cartoons in red: U-insertion RNAs. Cartoons in blue: U-deletion RNAs. $T_m$-values are in degree Celsius and are derived from ≥2 UV-melting measurement with SD-values ≤2%. Fluorophore positions are shown as stars. Yellow dots mark PS-modifications.

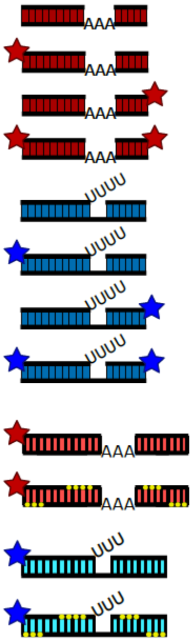


| RNA-substrate designation | $T_m$ (°C) 1st trans. | $T_m$ (°C) 2nd trans. |
|---|---|---|
| Standard U-insertion | 47.0 | 68.0 |
| | 47.0 | 68.0 |
| | 46.5 | 67.3 |
| | 47.0 | 68.0 |
| Standard U-deletion | 55.0 | 64.0 |
| | 55.0 | 63.7 |
| | 55.3 | 63.6 |
| | 56.0 | 64.5 |
| Non-natural U-insertion | 43.0 | 65.0 |
| Non-natural, PS-modified U-insertion | 41.0 | 62.0 |
| Non-natural U-insertion | 39.0 | 65.0 |
| Non-natural, PS-modified U-insertion | 41.1 | 64.1 |
